# Conserved Aberrant Developmental Trajectories of Human and Mouse SBMA Motor Neurons

**DOI:** 10.1101/2025.09.17.674754

**Authors:** Helen Devine, Martha J Roberts, Oliver J Ziff, Michael G Hanna, Linda Greensmith, Rickie Patani, Bilal Malik

## Abstract

Spinal bulbar muscular atrophy (SBMA) is a neuromuscular disease caused by a polyglutamine repeat expansion in the androgen receptor gene (AR). Lower motor neuron loss is a key feature of the disease, yet it remains poorly understood why these cells are affected. The transcriptional mechanisms underlying SBMA pathogenesis and how these evolve across developmental and disease stages remains incompletely defined.

To elucidate the molecular mechanisms underlying motor neuron loss in SBMA, we first performed transcriptomic profiling of both induced pluripotent stem cell derived motor neurons (iPSC-MNs) generated from SBMA patients and laser-captured micro dissected motor neurons (LCM-MNs) from symptomatic AR100 SBMA mice. We compared differential gene expression between the two models to identify shared transcriptional programs. To address the temporal progression of molecular changes we conducted profiling at key stages of motor neurogenesis in the developing iPSC-MNs and at pre-symptomatic and end-stage disease in AR100 SBMA mice to elucidate the emergence of the transcriptional phenotype and the trajectory of the gene expression changes.

We found significant transcriptional convergence between these two species. Notably, shared dysregulation was observed in pathways related to the spliceosome, the cell cycle and mitochondrial function. These transcriptional alterations emerged early in motor neurogenesis suggesting a developmental component to SBMA. Further in AR100 LCM-MNs we also observed disruption of mitochondrial and DNA damage repair pathways from pre-symptomatic to end stage disease.

This study identifies conserved pathogenic mechanisms across two SBMA model systems and provides crucial insights into the molecular basis and temporal dynamics of SBMA progression which may help identify potential therapeutic targets for SBMA.

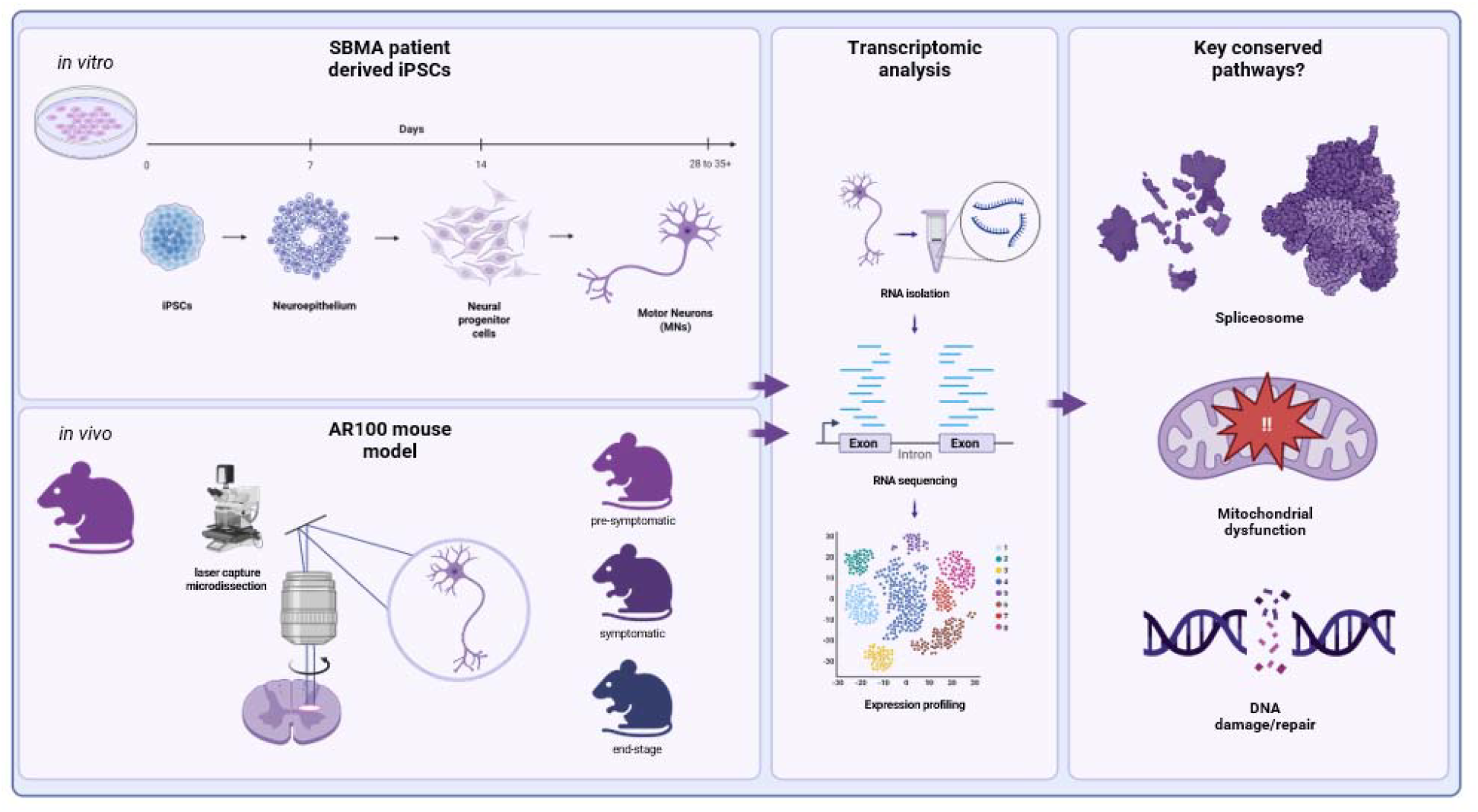

## Introduction

Spinal bulbar muscular atrophy (SBMA), also known as Kennedy’s disease, is an X-linked, slowly progressive neuromuscular disease which presents with neurological (lower motor neurons (MNs) and muscles) as well as systemic (metabolic syndrome, androgen insufficiency and neuro-urological) features^1^. It is caused by a CAG trinucleotide polyglutamine (polyQ) expansion in exon 1 of the androgen receptor (AR) gene on the long arm of the X chromosome^2^. Whilst the underlying causative genetic mutation was identified in 1991, there are still no effective disease-modifying treatments for this rare neurodegenerative disease. A major barrier to drug discovery in SBMA is the limited understanding of the molecular basis of disease^3^. Further it remains unclear which pathological processes are causally implicated in disease pathogenesis, and which arise secondary to neurodegeneration^4^.

Transcriptional analysis is a powerful tool to investigate the molecular mechanisms of disease by enabling systematic identification of dysregulated genes and pathways. Over recent years human stem cell-derived motor neuron (iPSC-MN) models have been instrumental in elucidating disease mechanisms in neurodegenerative disorders such as amyotrophic lateral sclerosis (ALS)^5–7^ and, more recently, in SBMA^8,9^. Patient-derived iPSC models offer the advantage of cell-type specific, human-specific context and genetic fidelity. However, iPSC-MNs are limited by being at a developmental stage of neuronal maturation which is a limitation for the study of progressive, adult-onset neurogenerative diseases such as SBMA. In this study, we complemented the iPSC-MN model approach with the well-characterised AR100 transgenic mouse model which expresses human polyQ AR with 100 repeats. This mouse model has been shown to have a clear SBMA-relevant neuromuscular phenotype including a progressive loss of motor neurons and AR aggregation. This model provides an *in vivo* context as well as allowing for a longitudinal study of the age-related aspects of disease. Using laser-capture microdissection we were able to isolate pure populations of MNs for transcriptomic analysis.

Combining transcriptional analyses from SBMA patient iPSC-MNs and LCM-MNs from symptomatic AR100 mice enabled us to perform a cross-species, temporally resolved analysis of gene expression dynamics in SBMA. This approach offers a unique opportunity to identify conserved transcriptional programs. Furthermore, by using a developmental iPSC-MN differentiation protocol we were able to capture early-stage transcriptional changes at key stages of motor neurogenesis that may prime MNs for later degeneration. We also isolated LCM-MNs from AR100 mice at pre-symptomatic, symptomatic and end-stage disease in order to examine the transcriptional changes in motor neurons over the disease progression spectrum.

In this study, we report convergent transcriptional dysregulation implicating mitochondrial dysfunction and impaired genomic stability across both models. The presence of significant transcriptional convergence between these two species reinforces their pathogenic relevance. Pathways involved in the structural integrity of the motor neuron and genomic instability present at a developmental stage and mitochondrial dysfunction and enrichment of mRNA processing pathways were present throughout all disease time points in the AR100 LCM-MNs. Together, these analyses refine our understanding of the early and progressive changes of SBMA molecular pathology and support the use of integrated model systems to provide a foundation for targeted therapeutic strategies.

## Results

### 1. SBMA iPSC-derived motor neurons exhibit transcriptional dysregulation in DNA damage response, cell cycle, and mitochondrial pathways

To investigate cell-autonomous transcriptional changes associated with SBMA we first generated highly enriched spinal iPSC-MN cultures from SBMA patients and unaffected controls (**Figure 1A and 1B**). There was no difference in the ability to terminally differentiate MNs between SBMA and control lines (**Figure 1C**). Reflecting the fact that SBMA is a slowly progressive neurodegenerative disease, iPSC-MN viability was not reduced using the Invitrogen LIVE/DEAD assay (**Figure 1D**). However, the Alamar Blue assay which is used to determine cell viability by determining which living cells are metabolically active demonstrated a significant reduction in metabolically viable cells (**Figure 1E**). We also examined the neurite length, neurite branching point and cell body clusters of the iPSC-MNs. SBMA neurite length was significantly reduced compared with controls, demonstrating impaired axonal growth which may indicate cytoskeletal instability or energy deficits (**Figure 1F**). Additionally, there were reduced neurite branching points in SBMA iPSC-MNs suggesting defective neuronal arborisation which may reflect altered network integration and synaptic function (**Figure 1G**). There was also reduced cell body cluster area which can be related to reduced survival, neurite loss or impaired cell-cell adhesion (**Figure 1H**). These findings demonstrate that the SBMA iPSC-MNs have a motor neuron phenotype.

**Figure 1:**
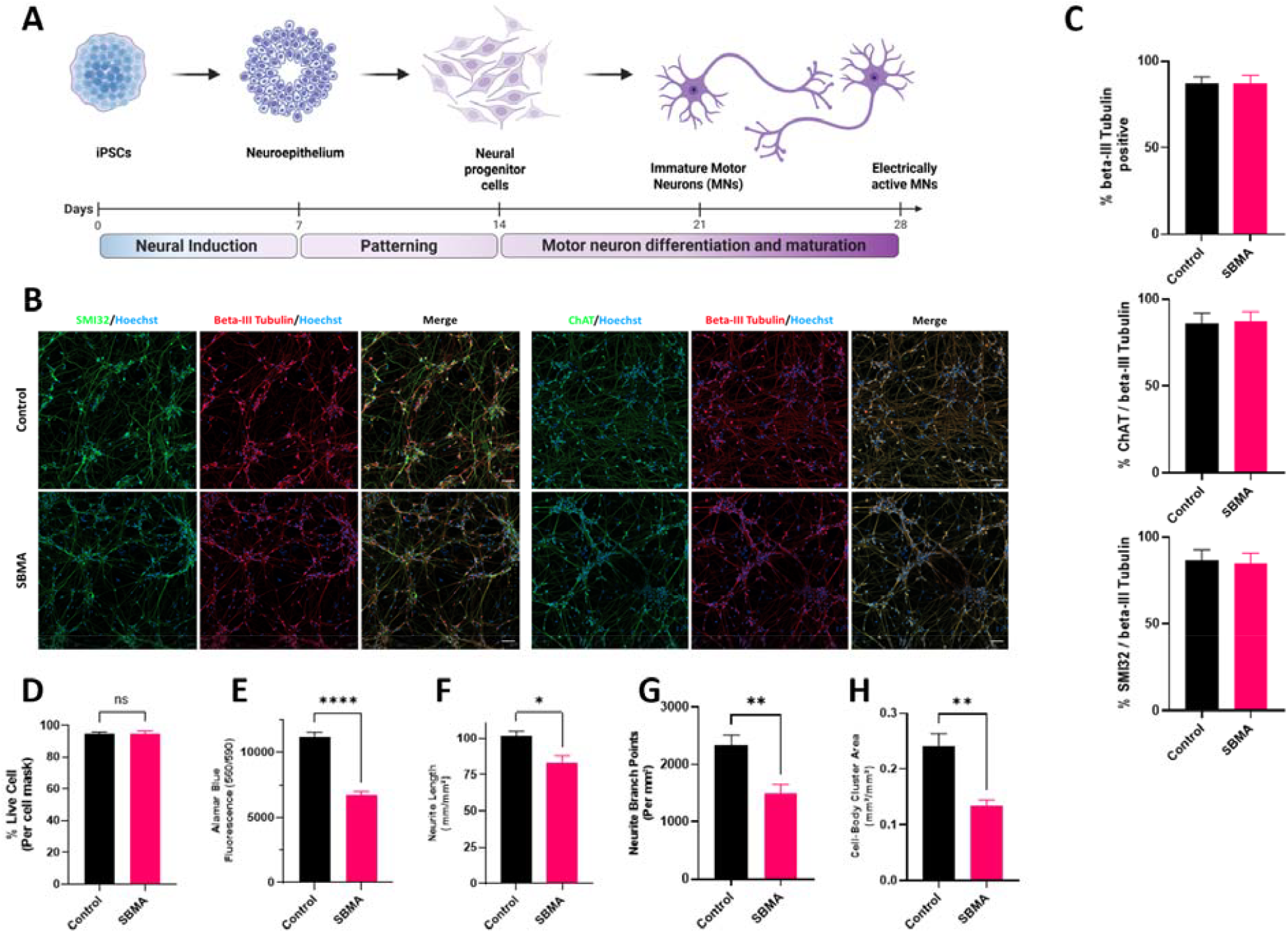
SBMA iPSC-derived motor neurons exhibit reduced viability and impaired neurite morphology despite normal differentiation efficiency. (A) Schematic depicting neurogenesis. Lines indicate sampling time-points in days. iPSC were obtained from four patients with confirmed SBMA, and four different healthy controls. One induction from each line was used for RNA-Seq. Induced pluripotent stem cells (iPSCs); neural precursor cells (NPCs); post-mitotic but immature motor neurons (d21); post-mitotic mature MNs (d28). Neural induction for 7 days with Dual SMAD and GSKb inhibitors; patterning for seven days with retinoic acid (RA) and sonic hedgehog (SHH); three days of low SHH, followed by NOTCH antagonist for coordinated cell cycle exit. (B) Representative immunocytochemistry images of iPSC-MNs with neuronal marker β-tubulin III to identify all neurons and motor neuron markers SMI32 and ChAT to immunolabel motor neurons. Scale bar is 50µm. (C) Graphs show the efficiency of neuronal differentiation was not significantly different between SBMA and control lines using the proportion of β-tubulin III neurons over total cells. To compare motor neuron specific efficiency the relative proportion of MNs (ChAT and SMI32 cells) to all neurons (i.e. of all β-tubulin III cells) was used and shows no significant difference between SBMA and control lines. Data is expressed as mean ± SEM from 4 SBMA and 4 control lines. (D) SBMA iPSC-MN was not reduced compared to control iPSC-MN however the number of metabolically viable cells was significantly reduced (E). Neurite length (F), neurite branch point (G) and cell-body clusters were all significantly reduced. Data expressed as mean ± SEM from 4 SBMA and 4 control lines. (*⍰= ⍰p ⍰< ⍰0.05, ** ⍰= ⍰p ⍰< ⍰0.01, *** ⍰= ⍰p ⍰< ⍰0.001, and **** ⍰= ⍰p ⍰< ⍰0.0001).

Next, we analysed gene expression using high-throughput, poly-A selected RNA-Seq derived from post-mitotic iPSC-MNs treated with dihydrotestosterone to elucidate the ligand-dependent activity of mutant PolyQ ARs. We performed differential gene expression analysis and found 3399 differentially expressed genes (DEGs) in SBMA versus control iPSC-MNs, with 1923 upregulated and 1476 downregulated (padj<0.05) (**Table S6**). We compared DEGs with previously reported AR responsive genes compiled from a search of the Gene Set Enrichment Analysis website from the Broad Institute (http://www.broad.mit.edu/gsea/). Here differential expression of *CCND1, CDK6, CENPN, GSR, MAK, MYL12A, PTPN21, SLC26A2, SPCS3, TMEM50A, XRCC5, DNAJB9, HERC3, IDI1, INSIG1, RAB4A, SPDEF* and *UAF* overlapped with known AR responsive genes. The proteins encoded by these genes are involved in the DNA damage repair response, the response to cellular stress, protein transport and the cell cycle. We performed GO analysis on all DEGs and found that the upregulated terms were related to the cell cycle and DNA repair as well as metabolic functions including protein binding and DNA and RNA metabolism (**Figure 2A**). Furthermore, there were notably down-regulated GO terms related to transport and mitochondria. Functional biological pathway over-representation analysis revealed enrichment in DNA damage response pathways (homologous recombination, FDR=6.3×10^−4^; Fanconi anaemia, FDR=1.9×10^−3^; DNA replication, FDR=0.26) as well as cellular senescence (FDR=8.9×10^−4^; *MTOR, CCNB1, RB1*) and cell cycle (FDR=3.5×10^−5^; *TP53, CDK4, CHEK1*) pathways in SBMA relative to control d28T iPSC-MNs (**Figure 2B, Table S12**).

**Figure 2:**
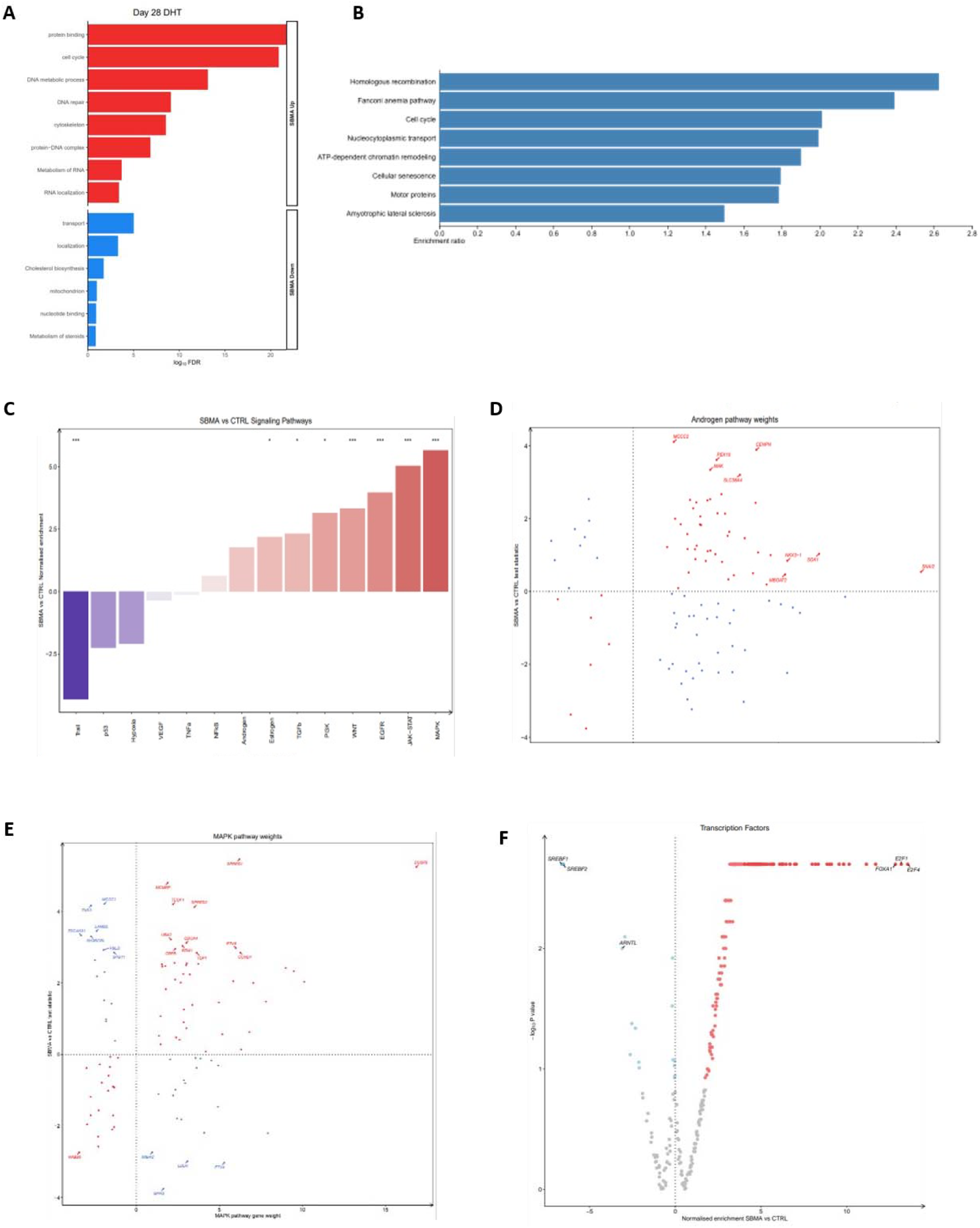
Transcriptomic and signalling pathway alterations in DHT-treated SBMA iPSC-derived motor neurons at day 28 of differentiation. (A) Functionally over-represented terms in up-regulated (red) and down-regulated (blue) differentially expressed genes using the hypergeometric test in DHT-treated SBMA versus control iPSC-MNs at day 28. (B) Enrichment analysis of the functional over-representation of KEGG pathways between DHT-treated SBMA and control iPSC-MNs at day 28 of motor neurogenesis, ordered by normal enrichment scores. (C) PROGENy signaling pathway activities in DHT-treated SBMA versus control using the weighted mean method. Pathways increased in SBMA are red and pathways decreased are blue. *** represents p<0.0001 and * represents p<0.05 (MAPK, JAK-STAT, EGFR, WNT, P13K, TRAIL p<0.0001; TGFb, Estrogen and Androgen p<0.05) (D) Expression changes of Androgen pathway signaling genes in SBMA versus control according to their PROGENy weights. Genes increasing androgen activity in SBMA are in red whilst genes decreasing androgen activity in SBMA are blue. (E) Expression changes of MAPK pathway signaling genes in SBMA versus control according to their PROGENy weights. Genes increasing MAPK activity in SBMA are in red whilst genes decreasing MAPK activity in SBMA are in blue. (F) Activities of transcription factors in DoRothEA inferred from the regulon expression changes in SBMA versus control. The normalised enrichment score in SBMA versus control (x-axis) is plotted according to the enrichment test p-value (y-axis).

Next, we performed Signalling Pathway RespOnsive GENes (PROGENy) analysis to further understand how signaling pathways are activated in SBMA iPSC-MNs^28^. This infers pathway activity changes based on perturbation experiments and weights genes based on responsiveness (**Figure 2C**). The most substantial pathway activity increase was in Mitogen-Activated Protein Kinase (MAPK; NES+5.66, p<0.001), followed by JAK-STAT (NES+5.03, p<0.001), Epidermal Growth Factor (EGFR; NES+3.96, p<0.001) and Wnt (NES+3.32, p<0.001). The most substantial decrease was in Trail (NES-4.31, p<0.001). As SBMA is caused by a mutation in the *AR* gene, we examined each gene in the androgen pathway and found the greatest responsiveness increases in SBMA iPSC-MNs in *CENPN, PEX10, MAK, SGK1* and *SNAI2*. Interestingly, the genes most upregulated in the androgen pathway are primarily associated with cellular stress response, DNA damage repair, and cell cycle regulation. We also examined the genes in the MAPK pathway as this had the most substantial increase and found the greatest responsiveness increases in *DUSP6, SPRED1, SPRED2, MCMBP* and *CCND1*.

Finally, we inferred the activities of 429 transcription factors (TFs) from examining their regulon (a set of genes regulated as unit, under the control of a regulatory gene) expression within the DoRothEA database^28^. This demonstrated that the AR transcription factor was amongst those with the greatest increase in activity in SBMA neurons (NES+4.06, p⍰< ⍰0.001). Interrogating individual genes constituting the AR TF regulon revealed the greatest increases in SBMA iPSC-MNs in BABAM2, FAM107B, SUOX, CDC6, PDZRN3, TNS3, and *MED1*. The proteins encoded by these genes are involved in processes including the cell cycle, mitochondrial function and cell migration. The TFs most increased in SBMA included E2F4, E2F1 and FOA1 whilst the strongest TF decreases in SBMA were in SREBF1, SREBF2 and ARNTL (**Figure 2F**). These TFs are related to cell cycle activity, enhancing neurite outgrowth and cholesterol pathways. Overall, these unbiased analyses of signal pathways and transcription factor activity converge on a cellular state defined by heightened stress responses, aberrant cell cycle signaling, and altered metabolism.

### 2. AR100 LCM motor neurons show altered metabolic, synaptic and neurodegenerative gene expression

To investigate transcriptional changes *in vivo*, we performed RNA-seq on laser capture micro dissected (LCM) spinal cord motor neurons (LCM-MNs) from symptomatic AR100 and AR20 (control non-pathogenic transgenic) mice at 12 months. We identified 9,817 DEGs (padj<0.05) between the AR100 and AR20 LCM-MNs (**Table S8**). Functional biological pathway over-representation analysis (**Figure 3A**) revealed enrichment in pathways of neurodegeneration (FDR=2.79e-8) as well as other neurodegenerative diseases such as Alzheimer’s disease (FDR=0.00002) and amyotrophic lateral sclerosis (FDR= 0.00037245). There is also enrichment of metabolic pathways (FDR=0.00001), endocytosis (FDR=0.0004) and axon guidance (FDR=0.042) (**Table S14**). These findings highlight the critical role of neurodegenerative pathways, metabolic and synaptic processes in SBMA LCM-MNs.

**Figure 3:**
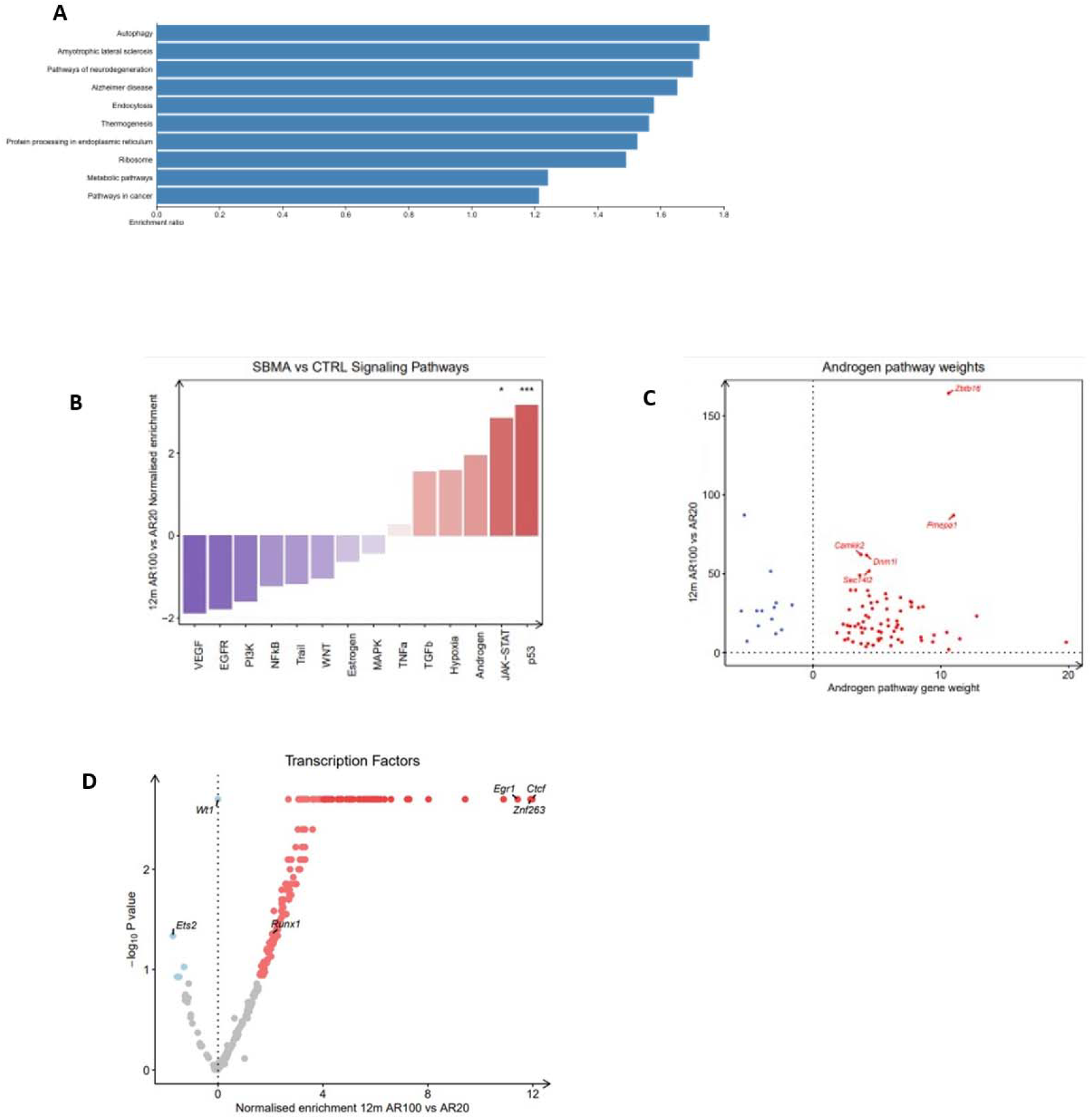
Symptomatic AR100 LCM motor neurons show altered metabolic, synaptic and neurodegenerative gene expression. (A) Enrichment analysis of the functional over-representation of KEGG pathways between AR100 and AR20 mice at 12 months, ordered by normal enrichment scores. (B) PROGENy signaling pathway activities in AR100 versus AR20 mice using the weighted mean method. Pathways increased in SBMA are red and pathways decreased are blue. *** represents p<0.0001 and * represents p<0.05. (C) Expression changes of Androgen pathway signaling genes in SBMA versus control according to their PROGENy weights. Genes increasing androgen activity in SBMA are in red whilst genes decreasing androgen activity in SBMA are blue. (D) Activities of transcription factors in DoRothEA inferred from the regulon expression changes in SBMA versus control. The normalised enrichment score in SBMA AR100 versus control AR20 (x-axis) is plotted according to the enrichment test p-value (y-axis).

We also performed PROGENy analysis comparing AR100 and AR20 mice at the 12-month symptomatic time point (**Figure 3B**). This revealed that the most substantial pathway activity increased was in the p53 pathway (NES 3.2, p <0.001) and JAK-STAT (NES 2.9, p=0.02). Again, we examined each gene in the androgen pathway (**Figure 3C**) and found the greatest responsiveness increases in AR100 LCM-MNs in *Zbtb16, Pmepa1, Camkk2, Dnm1l* and *Sec1412*. These genes are involved in cell cycle progression, neurite outgrowth and synapse formation, mitochondrial fission and biogenesis of Golgi-derived transport vesicles respectively. We then determined transcription factor activity using the DoRothEA database (**Figure 3D**). Here we found that the transcription factors most increased in AR100 LCM-MNs included *Ctcf, Znf263* and *Egr1* which are involved in chromatin binding, DNA-binding and mitogenesis. Together these findings show that SBMA motor neurons display transcriptional alterations across neurodegenerative and metabolic pathways.

### 3. Shared transcriptional alterations in RNA processing, cellular ageing, and metabolism define SBMA motor neurons across species

Having investigated the transcriptional changes in the iPSC-MNs and LCM-MNs in isolation, we next aimed to compare conserved and divergent disease signatures between the SBMA iPSC-MN and SBMA AR100 LCMs as these might help to identify the most impactful therapeutic targets. We first performed mapping of differentially expressed mouse genes from the 12-month timepoint dataset to their human orthologs (assuming conserved functions), based on Ensembl BioMart data, to ensure high-confidence gene pairs. Next, we DEGs in AR100 with DEGs in IPSC-MNs and undertook a pathway enrichment analysis. The goal was to identify biological pathways significantly represented in the homologous gene set, providing insights into conserved molecular functions and processes.

Of 9817 total mouse genes we successfully mapped 85% to human orthologs with 15% predicted to be species-specific genes (e.g., Gm49660). The mapped human gene symbols were compared to the human iPSC-MN dataset. A total of 1,125 genes were identified as common between the datasets and used for conserved pathway analysis. This comparative analysis revealed enriched pathways of the overlapping genes between human iPSC-MNs and AR100 symptomatic LCM-MNs and includes pathways related to neurodegenerative diseases and ALS as well as pathways related to mechanisms of neurodegeneration including nucleocytoplasmic transport, cellular senescence, autophagy and metabolic processes. The most enriched pathway is that of the spliceosome (**Table 1**, fold enrichment 3.4, FDR = 8.0E-05). The intrinsic vulnerability of neurons to splicing defects positions alternative splicing as a potential driver of SBMA pathogenesis, echoing its established role as a molecular hallmark of ALS^10^. Here, we found that there are common enriched pathways between SBMA human and mouse models at symptomatic time points with spliceosome dysfunction emerging as the most enriched pathway, alongside other mechanisms implicated in neurodegeneration including autophagy, nucleocytoplasmic transport and shared metabolic dysregulation.

**Table 1:** Overlapping transcriptional signatures in human iPSC-MNs and AR100 motor neurons highlight convergent disease pathways.

| Pathways | nGenes | Pathway Genes | Fold Enrichment | FDR |
| --- | --- | --- | --- | --- |
| Spliceosome | 22 | 132 | 3.4 | 8.0E-05 |
| Nucleocytoplasmic transport | 18 | 108 | 3.4 | 5.7E-04 |
| Ribosome biogenesis in eukaryotes | 11 | 77 | 2.9 | 4.7E-02 |
| Amyotrophic lateral sclerosis | 47 | 363 | 2.6 | 5.1E-07 |
| Cellular senescence | 20 | 156 | 2.6 | 3.9E-03 |
| Autophagy | 16 | 141 | 2.3 | 4.7E-02 |
| Salmonella infection | 28 | 249 | 2.3 | 2.8E-03 |
| Tight junction | 19 | 169 | 2.3 | 2.8E-02 |
| Pathways of neurodegeneration | 46 | 475 | 2 | 8.8E-04 |
| Endocytosis | 24 | 252 | 1.9 | 4.7E-02 |
| Metabolic pathways | 109 | 1527 | 1.4 | 3.3E-03 |

### 4. Progressive Emergence of a Transcriptional Phenotype During Motor Neurogenesis in SBMA iPSC-MNs

Having identified conserved pathways between human and mouse models of SBMA we then aimed to determine when these pathways emerge. We therefore analysed gene expression using high-throughput, poly-A selected RNA-Seq derived from the following developmentally categorized phases of lineage restriction: iPSCs (d0), post neural induction (d7), post patterning to the pMN domain (d14), post-mitotic but immature iPSC-MNs (d21). Initial principal component analysis identified separation along PC1 and PC2, corresponding to stages in motor neurogenesis. Once established as MNs, there was not clear separation between ‘immature’ and ‘mature’ MNs (**Figure 4A**). We performed differential gene expression analysis at each stage of motor neurogenesis to investigate the emergence of a transcriptional phenotype. We found an overall increase in the number of differentially expressed genes (DEGs) through differentiation into a post-mitotic MN fate. In iPSCs, there were 137 DEGs (110 increased, 27 decreased, **Table S1**) (false discovery rate (FDR) <0.05), at d7 there were 151 DEGs (105 increased, 46 decreased, **Table S2**), at d14 there were 22 DEGs (10 increased, 12 decreased, **Table S3**). As immature iPSC-derived MNs (iPSC-MNs) at d21, 395 genes (279 increased, 116 decreased, Table S4) were differentially expressed, and as established iPSC-MNs at day 28 we found that there were 477 DEGs between SBMA and control (246 increased, 231 decreased, FDR <0.05, **Table S5**).

**Figure 4:**
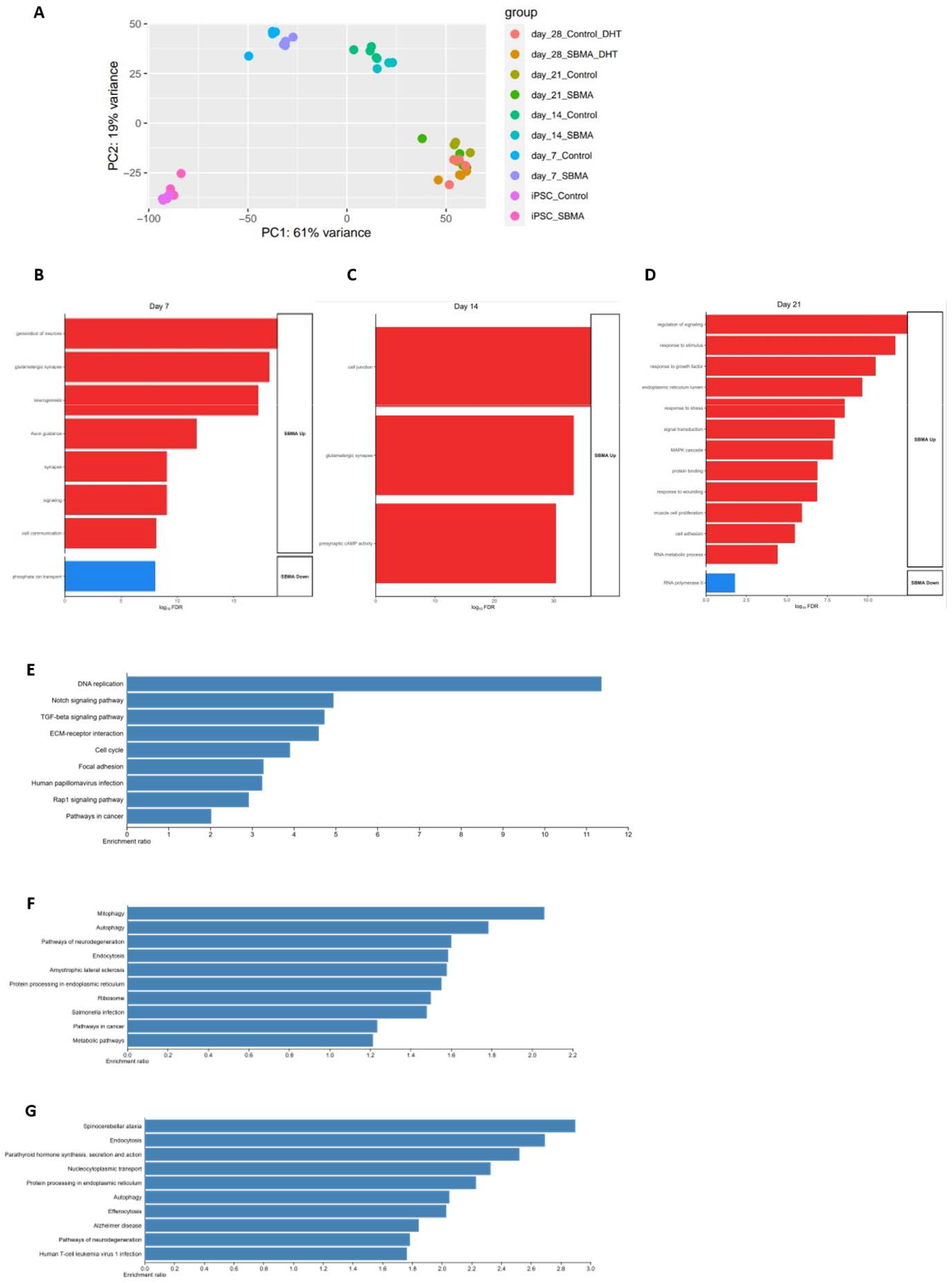
Disruption of morphological and DNA repair pathways emerges during early motor neuron development in the SBMA iPSC model. (A) Principle component analysis shows different time points are distinct populations and discriminates between SBMA and control lines. Individual cell lines demonstrate clustering at different developmental time points iPSC, d7, d14. There is less distinction between post-mitotic MNs at d21 and d28. No individual line is a consistent outlier. (n=4 control, n=4 SBMA) (B) (C) and (D) Functionally over-represented terms in up-regulated (red) and down-regulated (blue) differentially expressed genes using the hypergeometric test in SBMA versus control iPSC-MNs at day 7 (B), day 14 (C) and day 21 (D). (E) Enrichment analysis of the functional over-representation of KEGG pathways between SBMA and control iPSC-MNs at day 21 of motor neurogenesis, ordered by normal enrichment scores. (F) and (G) Enrichment analysis of the functional over-representation of KEGG pathways between AR100 and AR20 LCM-MNs from presymptomatic (3 months) and late-stage (18m) mice, ordered by normal enrichment scores.

Next, we performed functional pathway over-representation analysis using WebGestalt and Gene Ontology (GO) analysis on differentially expressed genes at each time point. We did not identify any significant enrichment of pathways between SBMA and controls at the iPSC stage, however at d7 (**Figure 4B**) there was GO enrichment for neurogenesis, synapse and cell signalling terms (padj<0.05). There was no over-represented pathway enrichment in SBMA neural precursor cells at day 14, but there was an increase in GO terms for the cell junction, glutamatergic synapses and presynaptic cAMP activity at this time point (**Figure 4C**). In early MN formation (d21), functional over-representation analysis (**Figure 4E, Table S10**) revealed upregulated genes were enriched for DNA replication (FDR=2.1×10^−5^; *DNA2, LIG1, MCM3*), and the cell cycle (FDR=0.01; *CDC23, RBL1, WEE1*) as well as pathways implicated in the structural integrity of the developing MN including the extracellular matrix-receptor interaction (FDR=3.3×10^−3^; *COL4A1, FN1, ITGAV*), notch signalling (FDR=0.03; *DTX3L, NOTCH2, NOTCH3*) and adherens junctions (FDR=0.03; *ACTN1, ERBB2, TJP1*), with GO term enhancement which included response to stress, MAPK cascade as well as regulation of signalling and cell adhesion (**Figure 4D**). Together, these findings indicate that the transcriptional phenotype of SBMA emerges progressively during lineage restriction with early perturbations in neurodevelopmental and synaptic pathways followed by dysregulation of cell cycle, stress response and structural integrity processes as motor neurons mature.

### 5. Presymptomatic AR100 motor neurons show early activation of neurodegenerative and mitochondrial stress pathways

Having identified overlapping conserved enrichment of disease pathways in LCM-MNs from symptomatic AR100 mice and SBMA patient iPSC-MNs, we next investigated whether these pathways were also present at an earlier stage of disease in the AR100 mouse model. We performed RNA Seq on LCM-MNs from 3-month-old presymptomatic AR100 and AR20 mice and found that there were 9805 differentially expressed genes (padj<0.05), with 5129 upregulated and 4676 downregulated genes (**Table S7**). Functional overrepresentation analysis (KEGG) revealed enriched pathways which overlap with other neurodegenerative diseases including spinocerebellar ataxia and Huntington disease, which, like SBMA, are both polyglutamine repeat diseases, and ALS, as well as mitophagy, oxidative phosphorylation, metabolic, and ribosome and spliceosome pathways (**Figure 4F, Table S13**). These findings indicate that the dysregulated pathways observed at the symptomatic time point are already dysregulated early in disease, prior to symptom onset. Next, we investigated the transcriptional pattern in LCM-MNs from 18m old late-stage disease AR100 mice. There were 1806 DEGs (padj<0.05) with Interestingly, there was enrichment of several key pathways (KEGG) including endocytosis, nucleocytoplasmic transport and autophagy as well as related to neurodegenerative diseases including spinocerebellar ataxia and Alzheimer’s disease (**Figure 4G, Supplementary Table 15**). This shows that key neurodegenerative, metabolic and RNA processing pathways are already perturbed in presymptomatic AR100 motor neurons and evolve to encompass additional processes such as autophagy, endocytosis and nucleocytoplasmic transport at late disease stages.

## Discussion

In this study we systematically investigated the transcriptional changes associated with spinal and bulbar muscular atrophy (SBMA) by integrating two complementary model systems: human SBMA patient iPSC-derived motor neurons (iPSC-MNs) and laser-capture micro dissected motor neurons (LCM-MNs) from the AR100 mouse model. We found similar alterations in pathways associated with mitochondrial dysfunction, genomic instability and neurodegeneration thereby suggesting that core pathogenic mechanisms are preserved across both models. We also demonstrated dynamic alterations in the transcriptional profile at different developmental and disease stages, providing a novel insight into the temporal evolution of the transcriptional profile in SBMA.

A key finding of our study is the convergence of the transcriptional signature between the iPSC-MNs and the LCM-MNs from symptomatic AR100 mice despite species and model system differences. The gene expression profile overlapped between the two models, and this concordance reinforces the potential of iPSC-MN modelling for mechanistic studies and as a platform for drug screening in drug discovery for SBMA. An interesting finding was that the most significantly enriched pathway across both models was the spliceosome, which suggests that RNA processing may be a fundamental and early driver of disease. Aberrant splicing has been implicated in the pathogenesis of other motor neuron diseases including ALS^10–12^ and spinal muscular atrophy (SMA)^13,14^. Neurons are vulnerable to splicing alterations^15^ and the dysregulation of the spliceosome may lead to further transcriptomic instability.

In addition to this, we observed consistent dysregulation of mitochondrial pathways and DNA damage repair and re-entry into the cell cycle. These findings align with current evidence that mitochondrial metabolism and genomic instability are central features of SBMA pathogenesis^16–20^. Neurons have a high metabolic requirement, with the main source of energy coming from oxidative phosphorylation in the mitochondria. Mitochondrial function diminishes during normal ageing^21,22^ and early mitochondrial impairment has been linked to neuronal loss in multiple neurodegenerative disorders^23–27^. Maintenance of the mitochondrial membrane potential is a metabolically demanding process and reduced mitochondrial ATP production has been observed in SBMA iPSC-MNs^16^. Furthermore, impaired DDR pathways have been identified as a potential common mechanism in polyglutamine diseases, leading to the hypothesis that the expanded CAG repeats in post-mitotic neurons pose a specific challenge for the DDR machinery^28^. Although neurons are post-mitotic cells and have exited the cell cycle (G0), they rely on cell cycle re-entry to initiate a DDR^29,30^. Cell cycle re-entry markers have been associated with other neurodegenerative diseases including ALS^31^and Alzheimer’s disease^32^. These studies have found increased expression of cell cycle proteins in conjunction with cell loss, leading to the hypothesis that aberrant cell cycle activation leads to the initiation of apoptosis, resulting in cell loss and ultimately neurodegenerative disease. Taken together, the consistent upregulation of mitochondrial pathways and the DNA damage response observed in this study may reflect a conserved stress response to the chronic misfolded protein burden seen in SBMA, in other polyglutamine repeat disorders and in other neurodegenerative diseases.

We also observed dysregulation of crucial structural integrity pathways during motor neurogenesis, which is consistent with our previous findings in primary MNs from the AR100 mouse model of SBMA^18^. This suggests that SBMA motor neurons may be developmentally primed for neurodegeneration. Furthermore, dysregulation in focal adhesion, extra-cellular matrix receptor interaction and cell signalling pathways, which have a major impact on synaptic behaviour, suggests that widespread disruption of MN cell signalling is an early pathological phenotype. The observed impairment in axonal and cytoskeletal gene expression suggests that SBMA MNs may experience substantial difficulties in neurite extension and synaptic connectivity, which are critical for neuronal function and survival. Additionally, alterations in mitochondrial gene expression, indicative of bioenergetic deficits, combined with dysregulated cell cycle regulators, point towards abnormal neurogenesis. These disruptions are particularly important as they may compromise neuronal stability and viability^33^. Furthermore, the degradation of extracellular matrix interactions observed in our study may exacerbate these issues, emphasizing the multifaceted impact of SBMA on neuronal development.

Whilst this study has several strengths including cross-validation across model and species systems and investigation of the temporal phenotype of transcriptional changes there are several limitations. Firstly, whilst iPSC-MNs represent a valuable developmental model to study disease-relevant mechanisms in a human context, iPSCs revert to an embryonic epigenetic profile with reprogramming and therefore may not truly recapitulate the disease process of neurodegeneration. Additionally, whilst this study focused on transcriptional dysregulation in an enriched pool of spinal cord motor neurons, this does not necessarily reflect the complex nature of the *in vivo* environment and particularly interaction with skeletal muscle which is now recognised to play a primary role in the pathogenesis of SBMA. Utilising an iPSC-derived iPSC NMJ model of SBMA may help to resolve these issues. Future work to functionally validate the transcriptional dysregulation of the spliceosome, mitochondrial function and genomic instability as well as taking a broader multi-omic approach will extend our findings.

In summary, the results of this study provide compelling evidence that SBMA MNs undergo early transcriptional disruptions which are likely to contribute to their dysfunction and eventual degeneration. Our findings identify conserved, dysregulated pathways across species, some of which are conserved over time and are therefore likely to contribute to disease pathogenesis. These include dysregulation of the spliceosome, mitochondrial function, DNA damage response, and cell cycle regulation. Critically, these transcriptional changes emerge during early motor neurogenesis, providing the first evidence of a developmental component to SBMA pathogenesis. By integrating cross species and temporal analysis, this work establishes the foundation for identifying new therapeutic targets and biomarkers in SBMA. Our findings also support the rationale for early intervention as many pathways are dysregulated even at a developmental stage and certainly at the pre-symptomatic stage, at least in AR100 mice. Furthermore, our results support the utility of iPSC-MNs in future mechanistic studies in SBMA as well as for use in drug screening for potential therapeutic intervention. Therapeutic strategies targeting mitochondrial and metabolic function or the DNA damage response may have potential in SBMA. Our discovery of conserved, cross-species transcriptional dysregulation in SBMA motor neurons provides new insight into understanding disease onset and progression and identifies molecular pathways that may be amenable to targeted therapeutic interventions at the earliest stages of disease pathogenesis.

## Methods

### iPSC samples and motor neuron differentiation

Patient-derived iPSC lines in this study have been characterised and previously described ^34^. Details of iPSC lines are described in supplementary table. iPSCs were differentiated into motor neurons as previously described^7^, generating enriched populations of motor neurons. Briefly, iPSCs were maintained on Matrigel (Corning Life Sciences) with Essential 8 Medium media (Life technologies), cells were passaged using EDTA (Life technologies, 0.5mM). 100% confluent cells underwent neural induction of 7 days with chemically defined medium containing 1μM Dorsomorphin (Millipore), 2μM SB431542 (Tocris Bioscience) and 3.3μM CHIR99021 (Miltenyi Biotec), patterning with 0.5μM retinoic acid and 1μM purmorphamine for 7 days with a 4 day expansion in 0.1μM purmorphamine and finally plated out for terminal differentiation in the presence of compound E. iPSC-MNs were treated with 10 µM DHT (Sigma) on d21, d24 and d27 to model the ligand dependent nature of SBMA by allowing translocation of the AR into the nucleus (d28T). SBMA and control iPSC MNs without treatment with DHT were also collected at d28. Cell cultures were maintained at 5% carbon dioxide and at 37°C. Cells were harvested for RNA extraction at relevant time points. Neurite outgrowth was observed using an IncuCyte system. The lengths of neurites, number of neurite branching points and cell body clusters was collected and analysed with NeuroTrack software (Sartorius).

RNA extraction was performed using the Promega Maxwell RSC simplyRNA kit (including DNAase treatment) and the Maxwell RSC instrument. All RNA samples had values over >2.0 for the 260/280mm absorbance using NanoDrop nucleic acid testing as well as a RIN score >8. The libraries of mRNA-Seq were prepared using the polyA_KAPA_mRNA_Hyper_Prep kit (Roche, UK). The multiplexed libraries were sequenced as 100 bp paired-end strand-specific reads using the HiSeq 4000 sequencing machine covering 40 million reads per sample. Quality quantrol of fastq files was assessed using FastQC with quality metrics for each fastq file, fastq screen, a low pass screen of several reference genomes for confirming the data maps to the intended species and multiQC which is an aggregator for the QC metrics. Technical repeats sequenced on different sequencing runs from the same samples were concatenated for downstream analysis. The files were then aligned to the human reference genome (human genome assembly GR Ch38) using STAR v2.6.1^35^. To assign and count mapped reads the FeatureCounts package was used to quantify transcript expression^36^. To perform differential gene expression analysis DESeq2 was used which uses a generalised linear model to normalise the gene counts ^37^.

### Laser Capture Microdissection of Spinal Motor Neurons from AR100 mice

All mice were bred and maintained at the UCL Queen Square Institute of Neurology Biological Services (London UK). All procedures were conducted in accordance with the UK Animals (Scientific Procedures) Act 1986, under appropriate Home Office licencing and were approved by the Institutional Animal Welfare and Ethical Review Body (AWERB). Male transgenic mice expressing either 100 (AR100 pathogenic) or 20 (AR20, non-pathogenic) polyglutamine repeats in the androgen receptor (AR) gene were generated by crossing with wild-type C57BL/6J females. Genotyping was performed via PCR using previously established protocols. In line with the male-specific presentation of SBMA, only male offspring were included in the experimental analyses. Study design adhered to the 3Rs to ensure ethical animal use and minimisation of numbers, whilst maintaining statistical robustness. Animals were kept in a controlled environment with regulated temperature and humidity. They had unrestricted access to standard chow and water and were maintained on a 12-hour light/dark cycle. Unfixed spinal cords were dissected from non-perfused AR100 and littermate control age and sex-matched mice. The lumbar segment of the cord was embedded in Shandon Mount (Fisher Scientific) and stored at −80°C. 14μm cryosections were obtained on PEN Membrane Slides (Zeiss). Sections were briefly dried and then sequentially passed in 70% ethanol, 100% ethanol, 1%Cresyl Violet in 100% ethanol, 100% ethanol for 1 minute each. Sections were immediately stored at −80°C. Microdissection was performed <48 hours after slide preparation. A Zeiss PALM Microbeam Instrument was used and sections were catapulted in Adhesive Cap 500 collection tubes (Zeiss). Motor neurons were identified by location and diameter >30μm and approximately 250 were captured for each sample Qiagen RLT plus buffer was added to the cap and incubated at RT for 20 minutes. Tubes were then briefly centrifuged, frozen and stored at −80°C.

RNA quality were assessed using a PicoChip or NanoChip on an Agilent Bioanalyser, as well as the TapeStation 4150 (Agilent Technologies) electrophoresis platform. For RNA-seq, libraries were be generated from total RNA using the Illumina TruSeq RNA kit v2 (TA muscle) or SmartSeq RNA kit (LCM low RNA input), according to the manufacturer’s directions. Libraries were sequenced at the UCL Genomic facility on a HiSeq 4000. All FASTQ files were analysed using FastQC software. The files were then aligned to the mouse reference genome GRCm38) using STAR^35^. To assign and count mapped reads the FeatureCounts package was used to quantify transcript expression^36^.

### RNA Seq analysis

Differential gene expression(DEG) analysis was performed in R using the DESeq2 package, which models count data using a negative binomial distribution and performs shrinkage estimation of fold changes and dispersion^37^. To account for excess zero inflation, especially relevant in small sample or sparse data contexts, we applied ZINB-WaVE for normalisation and dimensionality reduction prior to DEG analysis^38^. The ZINB-WaVE model accounts under a zero-inflated negative binomial distribution and estimates latent factors that capture unwanted technical variation (sequencing depth, batch effects).

Over-representation analysis (ORA) was conducted using WebGestalt ^39^ (WEB-based Gene SeT AnaLysis Toolkit [https://www.webgestalt.org/], version 2023). Input consisted of significantly differentially expressed genes (adjusted p < 0.05). Functional enrichment was assessed across Gene Ontology (biological process, molecular function, cellular component), KEGG^40^, Reactome, WikiPathways, and PANTHER pathway databases. Statistical enrichment was evaluated using the hypergeometric test with Benjamini-Hochberg false discovery rate correction (FDR < 0.05)^41^. Redundant terms were minimized via hierarchical filtering, and key results were visualized as bar plots and enrichment networks exported directly from WebGestalt. Gene Ontology (GO) and pathway enrichment analyses were also performed using ShinyGO (v0.77) [http://bioinformatics.sdstate.edu/go/]. The input gene list consisted of significantly differentially expressed genes (adjusted p < 0.05). Enrichment was assessed for biological process (BP), molecular function (MF), and cellular component (CC) GO categories, as well as KEGG pathways, using the Mus musculus genome as background. False discovery rate (FDR) was used to correct for multiple testing. The decoupleR package^42^ was used to estimate PROGENy signalling pathway activities and DoRothEA TF regulon activities inferred from gene expression changes.

### Statistics using Prism

GraphPad Prism 8 was used to perform statistical analysis on the experiments (n=4 biological repeats unless otherwise stated) and to generate graphs. Independent variables in different groups were analysed by ANOVA with post-hoc comparisons. P values of <0.05 was accepted as statistical significance with p<0.05=*, p<0.01 =** and p<0.001 =***). Error bars represent mean ± SEM.

## Keywords and abbreviations

SBMA: Spinal bulbar muscular atrophy
AR: androgen receptor
polyQ: polyglutamine
iPSCs: induced pluripotent stem cells
MNs: motor neurons, mouse model, mitochondria
DDR: DNA damage response
ALS: amyotrophic lateral sclerosis
DHT: dihydrotestosterone, time-series, MultiRNAflow

## Acknowledgements

We thank Dr Kenneth Fischbeck and Dr Chris Grunseich for providing the SBMA iPSC used in this study and the patients who so kindly donated their fibroblasts. We thank other members of the Greensmith and Patani labs for their input. We thank the Francis Crick Institute scientific platforms, particularly the advanced sequencing STP.

## Funding

This work was funded through an MRC Clinical Research Training Fellowship to H.D. H.D. was also in receipt of the Terrence Morris Memorial Scholarship funded by The Amar-Frances and Foster-Jenkins Trust and was also supported by the UCL Queen Square Institue of Neurology Kennedy’s Disease Research Fund. R.P. holds a Lister Research Prize Fellowship and gratefully acknowledges the generous support of Steve Redgwell, Liane Iles, Challenging MND, the Motor Neurone Disease Association (grant no. Patani/Dec22/957-793), My Name’5 Doddie Foundation (grant no. MN5DF/2022/003), and Target ALS (grant no. BB-2024-C4-L4). MGH is part of the ICGNMD Consortium, funded by an MRC strategic award to establish an International Centre for Genomic Medicine in Neuromuscular Diseases (ICGNMD) MR/S005021/1. LG is the Graham Watts Senior Research Fellow, supported by Brain Research UK. BM is supported by Brain Research UK.

## Competing interests

The authors have no conflicts of interest

## Notes

### Competing Interest Statement

The authors have declared no competing interest.

